# Characterization of the Post-Mating Responses of *Drosophila hydei*, a species that lacks Sex-Peptide

**DOI:** 10.1101/2025.07.31.667665

**Authors:** Maxime Revel, Zeynep Yildirim, Lea Fabbro, Emi Nagoshi, Robert K Maeda

## Abstract

Female post-mating responses (PMRs) in *Drosophila melanogaster* rely on proteins made by the male that are transferred to the female upon mating to modify her behavior and physiology. Many of the most characterized responses are initiated by a single protein produced by the male accessory gland called the Sex Peptide (SP). Yet many *Drosophila* species, even those living in similar environments, do not have SP. As PMRs are thought to be mechanisms used to maximize reproductive success, we wondered if PMRs are similar in species living in the same environment and how PMRs might be achieved in species that lack SP. Here, we investigate the post-mating responses of *D. hydei. D. hydei* is a Drosophilid that lacks SP, but is often found cohabitating with *D. melanogaster.* We show that the PMR in *D. hydei* is markedly different from that found in *D. melanogaster*; *D. hydei* do not increase egg production, nor do they limit remating to additional males or change their diet after mating. Instead, *hydei* seem to display only subtle PMRs related to activity patterns, in a way distinct from those characterized in *D*. *melanogaster*. These changes seem to have impacted things like lifespan and sperm selection, and demonstrate the wide variety of choices that species can make and be successful in a given environment.

## Introduction

Species have each evolved reproductive strategies to effectively allow for their propagation in a particular ecological niche. Female post-mating responses (PMRs), where mating triggers changes in female physiology and behavior, are part of these strategies. The inducers of these PMRs are often found to be male seminal fluid products that are transferred to females during mating. This is true for the fruit fly *D. melanogaster*, where it has been shown that seminal fluid proteins made by the male accessory gland (MAG) trigger many of the most studied PMRs (1) (2) (3).

Among the many *Drosophila melanogaster* seminal fluid proteins (SFPs) that are transferred to the female, the protein Sex Peptide (SP) has been found to induce many PMRs, including: increasing egg production and laying, changing female receptivity to additional matings (4) (5), shifting the female circadian activity(6) (7), and shortening female lifespan (8) (9) While many of the most studied PMR effects in *Drosophila melanogaster* are induced by the SP pathway, it turns out that SP is not found in several branches of the *Drosophila* phylogenetic tree (10, 11). This begs the question of whether or not PMRs found in these species will be similar to those of *melanogaster* and how any PMRs existing in these species are induced.

At first glance, one might logically expect that certain PMRs would be found almost universally, even in the absence of SP. This is particularly true of PMRs that might economize the use of resources. For example, the induction of egg production and laying in mated female would seem to be highly valuable; increasing egg production only upon mating would seem to be an efficient strategy to prevent unnecessary egg production when males are not available for fertilization. Other examples might be less clear, like the reduction of female remating. While this is debatably advantageous for males to ensure the use of their sperm, it seems less advantageous for females who might want to have the ability to trade-up from a first sexual partner, leading to a potential case of sexual conflict (12). That being said, there are other known examples of organisms that share this PMR with *melanogaster* despite the absence of SP. For example, the ground beetle females (*Leptocarabus procerulus*) also are refractory to remating like in *D. melanogaster* (13). Thus, one might imagine that the more closely related species of the *Drosophila* genus, especially ones living in similar environments, might share many PMRs even in the absence of SP.

We have recently become interested in the species *D. hydei*, a species that is part of the *repleta* group (Siphlodora subgenus) that diverged from *D. melanogaster* (S*ophophora* subgenus) about 60 million years ago (14). Like *D. melanogaster, D. hydei* is a cosmopolitan species, often cohabitating with *D. melanogaster* in the wild (15)(16). This might suggest that similar evolutionary pressures were present on both species and therefore, that these two species might have evolved similar reproductive strategies. However, the little we know about *D. hydei* PMRs suggests that this is not correct. First, *D. hydei* males (along with other species closely related to *hydei*) have been selected to have unusually long sperm, often measuring over 2.3 cm in length (about 10X longer than those of *melanogaster*) (17). This odd sexual ornamentation seems to have had significant consequences on their reproductive strategy, being associated with a delay to the onset of male reproductive maturity (*hydei* males take approximately 10 days to reach sexual maturity vs 3-5 days for *melanogaster*) and limiting the number of sperm transferred to the female during mating (18). Perhaps due to this selection for long sperm, a second significant difference from *melanogaster* is the loss of the SP gene. Recent work has shown that although SP exists in the Siphlodora subgenus, *D. hydei* is part of the *repleta/nannoptera* radiation, a branch of Siphlodora that have lost the SP gene (10). Consistent with the absence of SP in *D. hydei*, at least one phenotype associated with SP activity in *D. melanogaster* is not present in *hydei; D. hydei* females tend to be willing to remate many times over a short span of time (19).

In order to further our understanding of PMRs in non-*melanogaster Drosophila* species, we decided to look more carefully at the response of *D. hydei* females to mating. Our investigations show that unlike *D. melanogaster*, *D. hydei* do not seem to produce more eggs after mating. Instead, *hydei* females seem to produce and lay an equal number of eggs over a three-day period regardless of mating status. Interestingly, the distribution of eggs laid does change over these three days, as mating triggers a burst of egg-laying that lasts for about 24h, followed by a period of reduced egg laying. In *melanogaster*, mating has also been shown to reduce female lifespan. Numerous factors have been suggested to be the cause for this reduction of lifespan, from direct signaling from male seminal fluid components like SP, to the metabolic shift required for females to increase egg production (20). As *hydei* do not possess SP and lay a similar number of eggs regardless of mating status, it was notable that we found *hydei* females intermittently mated to males lived equally long lives as their virgin female siblings. Video-assisted activity monitoring was then used to examine the activity patterns of mated *hydei*. Using this method, we were able to uncover post-mating changes in the *D. hydei* female circadian activity as well as other behavioral changes. Lastly, we investigated sperm competition in *hydei* and how changes in accessory gland structure seem to modify sperm competitiveness. Overall, our results show that *D. hydei* seem to have adopted a very different, yet equally successful, reproductive strategy that allows them to coexist with species like *D. melanogaster* in the same environment.

## Results

### I – Mated females show a short-term egg laying burst with no increase in egg production in *D. hydei*

Most of the best characterized PMRs in insects have been studied in *D. melanogaster* where they are often linked to the seminal fluid protein SP. To investigate the post-mating responses in species without SP, we focused on *D. hydei,* a species that lives in similar environments, yet lacks SP. Consistent with the lack of SP in *D. hydei*, this species is known to lack the common SP-induced inhibition of remating; *D. hydei* females remate rapidly after an initial mating (19).

To identify possible post-mating responses of *Drosophila hydei*, we first investigated the female’s egg laying response. For this assay, we allowed individual females to mate freely over a two-hour period (generally mating one to three times) and then compared the egg-laying response of individual females over a three day period or longer. In *D. melanogaster*, female egg laying is increased over a period of up to 10 days (21). With *D. hydei*, we found that mated females only laid more eggs than virgin females for the first 24 hours post-mating (Fig. 1A and D.). During the subsequent days, mated females would, on average, lay less eggs than virgins, indicating the presence of a short-term response in ovipositioning. The total amount of eggs laid over three days post-mating was found to be identical between mated and virgin flies (Fig. 1B.), suggesting that the ovipositioning response in *D. hydei* is not accompanied by an increase in egg production. These results suggest that females may produce eggs at a constant rate, and that mating triggers a rapid release of stored eggs. After the short term response in post-mating egg laying, the egg laying behavior of mated females returns to a level identical to their virgin counterparts (Fig 1.D.) even though she will often continue to lay fertilized eggs.

**Figure 1.**
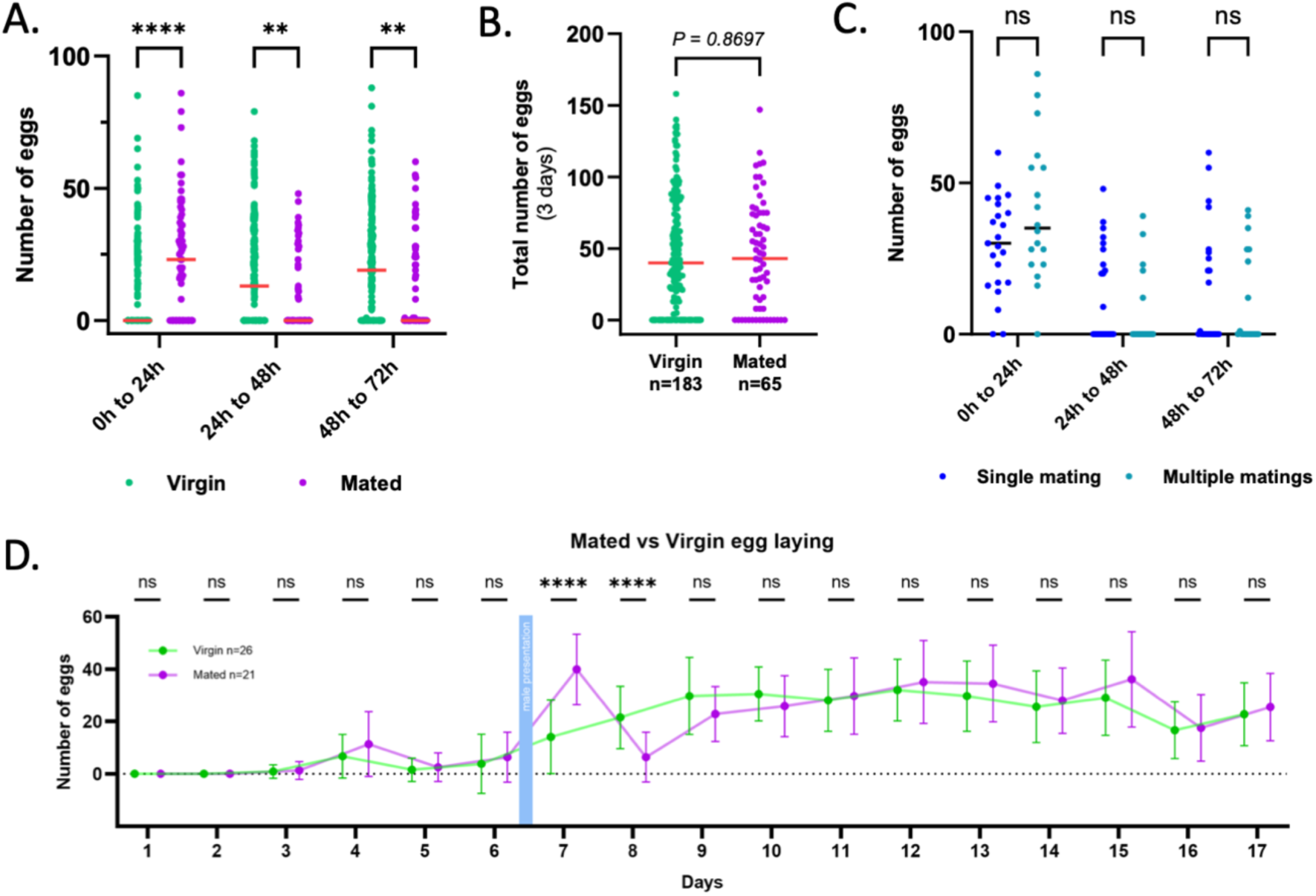
Egg laying in Drosophila hydei. A. Average number of eggs laid each day during the three days post-mating for females that have mated a single time, or more than once in a rapid succession. B. Total number of eggs laid by virgin or mated females for three days post-mating. C. Number of eggs laid by virgin or mated females in the three first days post-mating. D. Average number of eggs laid by individual females throughout the 17 days after eclosion. Flies represented by the green line were mated on day 6 to males aged for 10 days. The total amount of eggs for three days (B) was analyzed using a test of Mann-Whitney. Daily egg laying data (A, C. and D) were analyzed using a Mixed-effect model, for each day, conditions were compared using Šídák’s multiple comparisons test. ns (P > 0.05), * (P ≤ 0.01** (P ≤ 0.01), *** (P ≤ 0.001), **** (P ≤ 0.0001).

As *hydei* females mate multiple times during a typical mating assay, we sought to determine if the number of matings influenced the number of eggs laid. To do this, we compared the number of eggs laid by females that mated once, to those that mated twice or more times over the course of two hours. Looking at the first three days post-mating, we could not observe any significant difference between the two mating conditions (Fig. 1C). Thus, we find that rapid remating does not seem to result in the induction of a stronger egg-laying response.

### II - Study of the impact of egg-laying on the lifespan of females

Our egg laying assays show that unlike *D. melanogaster*, *D. hydei* females do not show a massive increase in egg production upon mating, but instead seem to produce eggs at a constant rate. In *melanogaster,* the induction of egg production is thought to help economize the energy females would use to produce eggs when males are not readily available. Phenotypically, this is thought to be partially responsible for the reduction of mated-female lifespan relative to males or virgin females (22, 23). We decided to ask if mating modifies lifespan in *hydei* where females seem to produce eggs, regardless of mating status, at a constant rate. To do this, we performed survival assays on both mated and virgin *hydei* females as well as virgin males. For these experiments, we housed flies in groups of approximately 10 flies per tube. For the mated female group, once a week, for three weeks (skipping the first week due to female immaturity), we added an equal number of fertile males and allowed mating overnight. Using this protocol, we find that virgin and mated females survive similar times (and less than their male siblings) (Figure 2A). Interestingly, in a separate experiment, we examined the survival of females and males housed together. In this experiment, females housed with males showed a significantly lower lifespan than virgin females. Thus, it seems that males or the act of constant mating may have a negative effect on *hydei* females that is not seen by intermittent matings (Supplementary Figure 1). This latter trend is similar to that observed in *D. melanogaster* where females undergoing intermittent matings lived longer than females continuously exposed to males (23)

**Figure 2:**
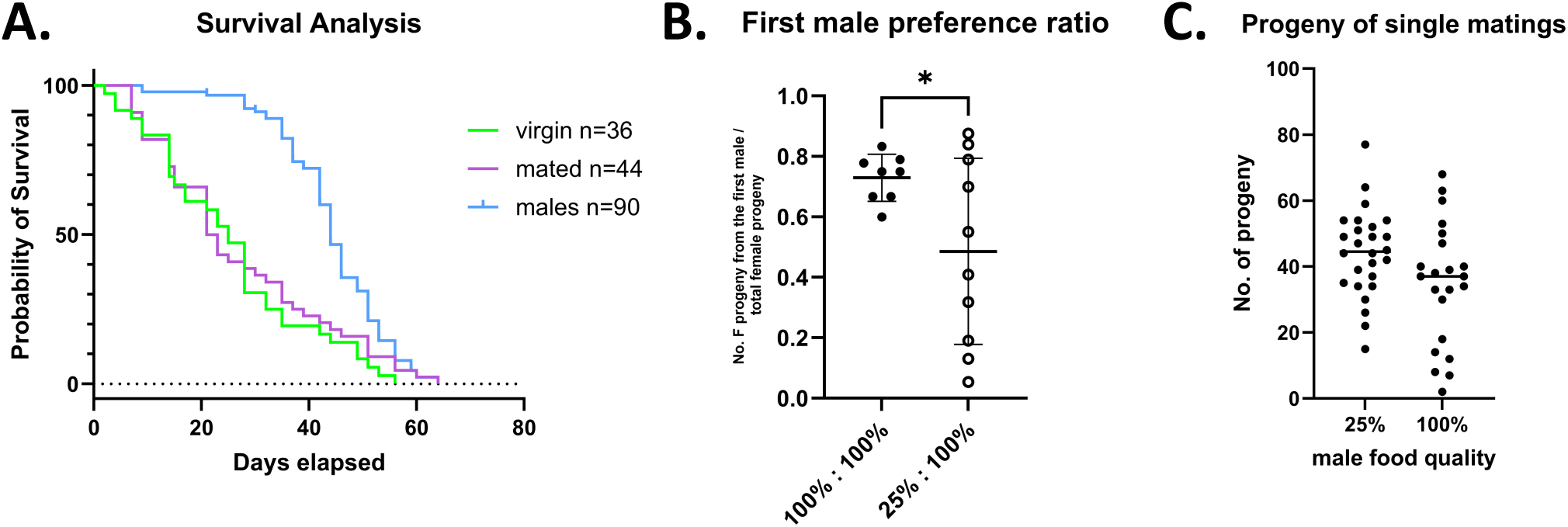
Potential impacts of constant egg-laying and multiple matings: A. Survival analysis of virgin females (green), mated females (magenta), and virgin males (blue) plotted as the percentage of total flies living at each day (Probability of Survival). Kaplan-Meier ( with Mantel-Cox and Gehan-Breslow Wilcoxon tests) analysis shows no significant difference between virgin and mated females but significant differences between both female populations and males (p<0.0001). B. Sperm competition assay between, males reared during all larval development on normal food (100%) or 25% diluted food (25%). Paternity of the female progeny could be determined by counting number of w- or w+ females. The growth conditions of the males are listed on the x-axis; the first male growth condition is listed before the colon and the second male growth condition is listed after the colon (* indicates that there is a significant difference between the two experimental set-ups, *p=*-0.0453, unpaired t-test). C. Total progeny from singly mated females from males grown under each growth condition. No significant difference between each condition was found.

### III - Male health and sperm competition

As mentioned above, we and others found that *hydei* females often mate multiple times during the timeframe of a mating assay. These matings are relatively short, generally lasting under 3 minutes. However, our data shows that multiple matings over this short timeframe does not seem to yield a statistical increase in the number of progeny produced (Fig. 1C). Thus, we wondered what advantage multiple matings would give to the species. One possible answer was that multiple matings might be a mechanism to allow female to select better sperm. In order to investigate this, we performed sperm competition assays by crossing *white* mutant *hydei* females to either *white* mutant males or *wt* males followed by immediately crossing these singly-mated females to second males of the alternate genotype (*w^+^* or *w^-^*). When the progeny of these crosses eclosed, we were able to score the eye color of the resulting female progeny to determine the male whose sperm was used by the female to fertize her eggs. In *melanogaster*, it is generally found that when secondary matings occur, there is a last male precedence, where the sperm of the last male copulating is used over that of previous matings (24). In *hydei*, we find the opposite to be mostly true, in that the first male’s sperm seems to produce more offspring than that of the subsequent male (Fig. 2B).

We then asked if this ratio could be modified based on different characteristics of the male. As our hypothesis was that this may be a method for the female to select sperm from healthier males, we sought out treatments that might subtly affect the male health. Previous studies have shown that the nutritional environment under which a male develops can effect male size and affects male fertility in Drosophilids (25). Interestingly, in *D. melanogaster*, the reduction in male fertility may be independent of the individual sperm cells themselves as sperm length seems to be similar between nutritionally minute flies and control flies (25).

In our sperm competition assay, we tested the ability of flies grown under reduced nutritional conditions (grown on 25% nutrient food) to defend against secondary sperm coming from a fly grown under standard conditions. Measuring the thorax size of flies grown under each condition confirmed that flies grown on 25% food were smaller than flies grown under standard conditions (supp fig 2A). To see if sperm size changed in these two populations, we then measured the length of the testes of flies grown under the two nutritional regimes (supp fig 2B). As testes length positively correlates to sperm length in flies and is much easier to measure than the tangled, multiple-centimeter long sperm of *hydei*, we, like others, decided to use this as a reasonable proxy for sperm length measurements (26). Based on the similarity of testes lengths, we believe that, like in *melanogaster*, sperm length is not affected by our manipulations (25). We then tested if a similar number of sperm was deposited by minute or wild type males. Previously, others have reported that minute males make and transfer less sperm than wild-type males. However, this seems to be visible through a cumulative effect from multiple matings from the undernourished male (27). Our results indicate that females mating once to either type of virgin male produces a similar number of offspring (Fig 2C), suggesting that a similar number of sperm are stored from males of each condition after a single mating to a virgin male. In competition, however, we found that sperm from nutritionally starved males (25% food) were less able to compete with the sperm of control males. As seen in Fig 2B, control males show a first male preference. However, this is reduced when the first male was raised on lower quality food. Overall, these results suggest that females may be able to determine male quality based on the seminal fluid that he deposits. Assuming the sperm itself is not changed (being of similar length in minute flies ((25) and supp figure 2B)) and that they are deposited in a similar amount, we hypothesize that the difference in sperm competition is due to either the female ejecting/manipulating the first male’s sperm based on external signals from the male (pheromones, size etc.) or from internal signals present in the two ejaculates.

### IV – The absence of SP has no direct consequences on the Male Accessory Gland shape or structure

Given the changes that we found in the sperm competition assay and the idea that differences in the seminal fluid may contribute to the loss of sperm competitiveness after being grown under less ideal circumstances, we decided to examine the effect of food reduction on MAG growth. We started this by first examining the MAGs of control flies from *melanogaster* and *hydei*. At the same time, we decided to examine the accessory glands of several *Drosophila* species from different parts of the Drosophila phylogenetic tree, selected based on availability, their phylogenetic position and their relationship to the SP gene.

In *D. melanogaster*, the MAG consists of a pair of sac-like lobes of around 800µm in length that are made up of two secretory cell-types. The main cells that form most of the gland and the secondary cells (SCs), which represent around 40-50 cells per lobe, that are located at the distal tip of the gland, interspersed with main cells. The SCs are larger than main cells and are filled with characteristic, large vacuole-like structures (28).

We first examined flies in the *Sophophora* subgenus. In this radiation, the SP gene has been shown to be highly conserved and has even undergone gene duplications (10). This subfamily is also characterized by the SP gene being translocated to a different location, near the gene *capricious* (*caps*), from its ancestral location, near the gene *NaPI-III* (11). For example, *D. melanogaster* has two copies of a SP-like gene near *caps*, while some species in the *ananassae* group have as many as 7 copies. Within the flies examined in the *Sophophora* subgenus, all had glands with relatively small lobes reaching no more than 1,000 µm in length (like in *Drosophila melanogaster* (Fig. 3A-B)). Furthermore, we found that the glands rarely contained more than 50 secondary cells per lobe, and that they were always located at the distal tip of the lobes (Figure 3A-B and Supp Figure 3). Interestingly, when examining the MAGs of non-*Sophophora* species (Fig. Fig 3A-B), glands tended to be longer, reaching up to 3,000 µm per lobe. Besides the length, we also found SCs spread all along the length of the lobes (Supp fig 3), rather than sequestered at one end. The number of SCs in these species was highly variable, from around 150 SCs per lobe in *D. busckii* (Fig. 3A-B.) to more than 400 in *D. nannoptera* and *up to* 500 in *hydei* (Fig. 3A-B). Interestingly, the dichotomy between species that have short glands with few SCs and those that have long glands with many SCs coincides with the divergence of the Sophophora group.

**Figure 3.**
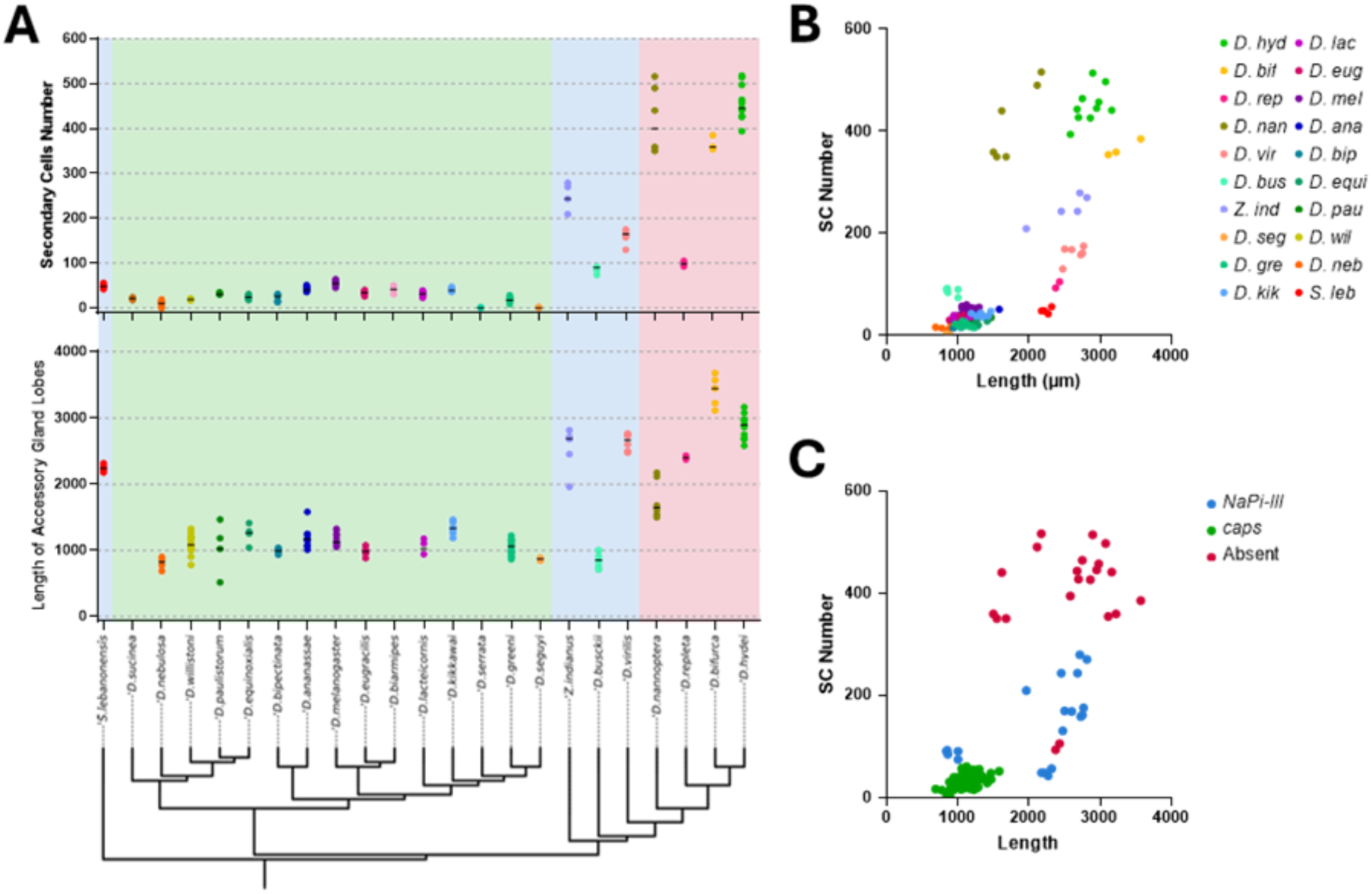
Accessory Gland lobe length and Secondary Cell numbers in Drosophila species. A. Phylogenic distribution of Secondary Cell number (top) and Lobe length (bottom). The Sophophora subgenus is highlighted in yellow on the phylogenetic tree.The location of the SP gene is indicated with background coloring: green – near caps; blue, near NaPi-III, red – absent. Phylogenetic tree based on (14). B. Number of Secondary Cells relative to the length of the lobes, colored per species. Species that are part of the Sophophora subgenus are highlighted in yellow. C. Number of Secondary Cells relative to the length of the lobes, colored in respect to the position of the SP gene.

To get an idea of the ancestral form of the gland, we decided to look at the glands of more basal species. Surprisingly, the glands of *Scaptodrosophila lebanonensis* are long, but with secondary cells located only in the center of the lobes (Supp fig 3.). Another species, *Chymomyza pararufithorax (*Supp. Fig. 4) also have long glands and have many SC spanning about ¾ of the gland from the distal tip. Based on these results, it seems likely that ancestral MAGs were probably more similar to what is found in the non*-Sophophora Drosophila*.

In *D. melanogaster,* the function the SCs seems to be tightly linked to the protein SP. Defects in the SCs leads to a phenotype where the PMR does not last for more than about 24 hours. This defect in the so-called “long-term PMR” is due to SP failing to be attached to sperm as they enter the sperm storage organ. We now know that SCs produce many of the proteins that attach SP to sperm allowing it to be slowly released over the following days to continually induce the SP response (29). As the *Chymomyza* lineage seems to be the first lineage where SP appeared (11), the finding of numerous SCs in a species of this group (*Chymomyza pararufithorax*) indicates that the presence of SP is not a prerequisite to having SCs and that SCs likely played a different role in male reproduction prior to its role in SP attachment. Meanwhile, the repeated loss of SP in *Drosophilids* that still have SCs, like *D. hydei* (10), indicates that these functions may still be present.

To investigate if our food manipulation might have the potential to change the qualitative composition of the seminal fluid, we examined the size and structure of the MAG after growth on different media concentrations. Previously, we used a single treatment condition where approximately 70-100 embryos were transferred into large tubes containing about 8 ml of food that was diluted to 25% of its normal nutrient concentrations. For a more precise measurement of the effect of the nutritional composition of our media on MAG structure, we placed precise numbers of embryos (*melanogaster* or *hydei*) into small tubes containing 3 ml of food diluted to four different food concentrations (100%, 50%, 25% and 10%). Upon eclosure, males were collected and transferred to normal food tubes to mature (five days for *melanogaster* and ten days for *hydei*). The thorax of some of these males was measured to confirm the effect of reduced nutrition on the adult flies. For some combinations of high egg density and low food concentrations, surviving adults could not be isolated. For this reason, we chose to dissect males from the 30 egg count density because adult flies could be obtained at all food concentrations. Dissection of these males showed that the accessory gland size and the numbers of SCs was affected by food quality, particularly at lower food concentrations (Fig. 4). Due to the link between MAGs and seminal fluid production, it seems likely that changes in male nutritional status could affect non-sperm constituents of the seminal fluid.

**Figure 4:**
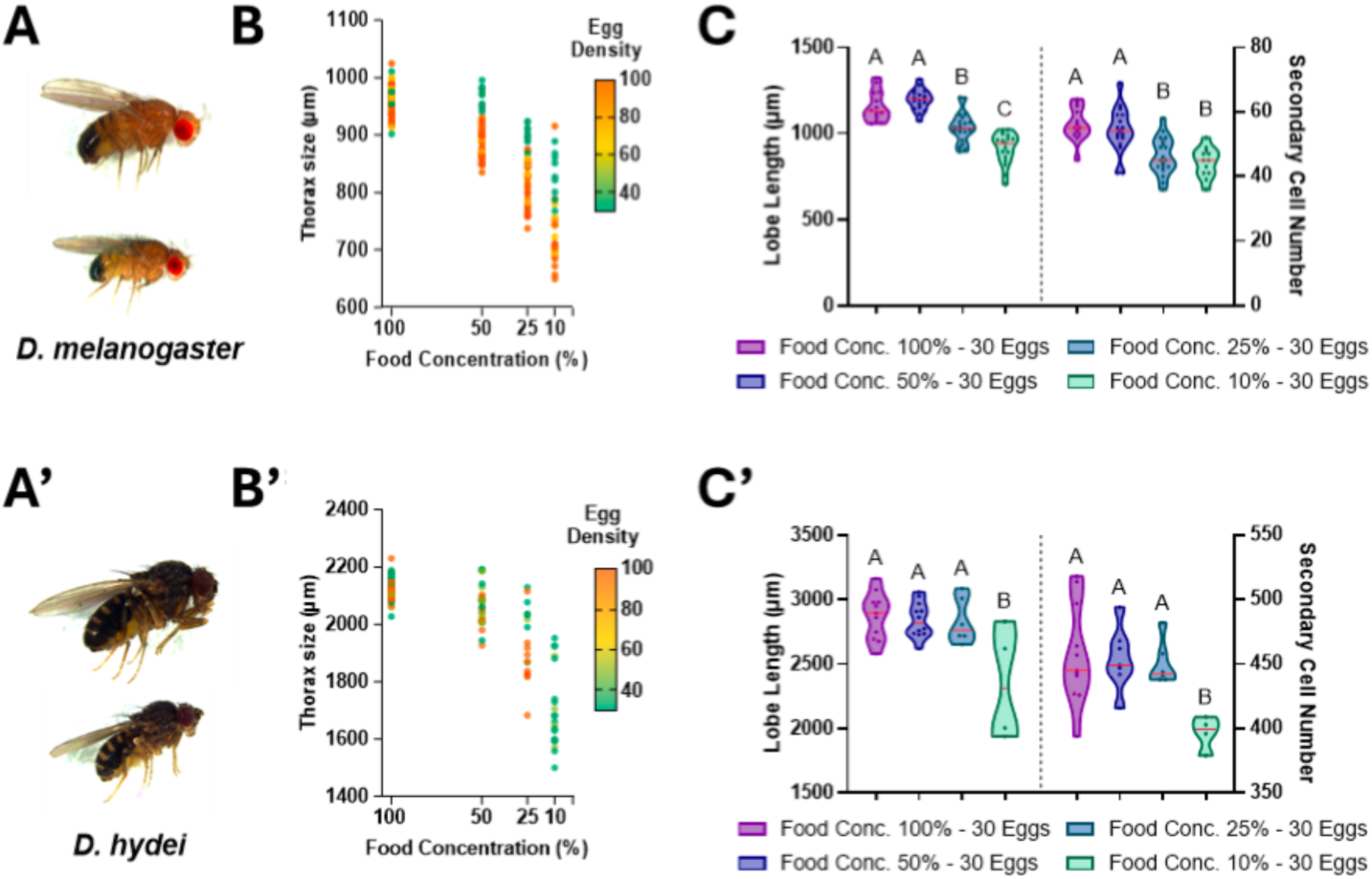
Thorax size and accessory gland morphology under starved developmental conditions. The species used were *D. melanogaster (top), and D. hydei (bottom*). A. and A’ Comparison of the flies’ size, at the same scale, between the most concentrated food (100%) and the least (10%). B. and B’ Thoracic size is represented as a ratio of the average size of the thoraces relative to the largest size of thoraces of flies reared on 100% food. The concentration of the food they’ve been reared on (x-axis), and the density of the eggs that were put in the tube (color). C. and C’ Measurement of the size of the gland’s lobe and the number of secondary cells they contain on different food concentrations. Analyzed using ordinary one-way ANOVA.

### V– Mating induces long-term changes in a highly stereotypical circadian activity

To uncover other post-mating responses in *hydei*, we decided to examine some of the known behavioral responses of *melanogaster* or other species. Previously, it has been reported that there is an increase in daytime locomotor activity in *D. melanogaster*, (siesta sleep, more pronounced in virgins) (*30*). Furthermore, it has been reported that there is a relative decrease in the activity of mated females just prior to circadian lights-on (morning anticipation, more pronounced in virgins) (6). Analyzing the locomotor activity of virgin *Drosophila hydei* females using the Drosophila Video-assisted Activity Monitor system (DrosoVAM)(31), we were able to document a highly stereotypical activity pattern in *D. hydei* with clear peaks of activity in the morning and evening (Fig. 5A). Although at a cursory glance, this pattern seems similar to that of *D. melanogaster*, it shows no sign of a classical morning anticipation. Indeed, *melanogaster* virgin females display an increase in activity as the time approaches circadian dawn/“lights-on”, while the activity of *D. hydei* increases only after the lights turns on. Under constant darkness conditions, these sharp peaks of activity during the day, particularly at “lights-on” slightly fade, indicating that it is mostly a startle response to the light (Supp fig 5A). Nevertheless, we do see an overall increase in activity during the day and an alternative kind of “morning anticipation”, characterized by a pre-dawn static period where activity drops for the last two to three hours before “lights-on” (Fig. 5 A-B). These activity trends are found in both virgin and mated females, as well as in males (Supp fig 5B). When examining the differences between mated and virgin females, we noticed that mated females seem to be less active overall but more noticeably during the nighttime hours. Averaging the distance moved for each hour of the day over three and a half days, we see that mated females move significantly less than virgins for a few hours in the middle of the daylight period and also for most of the night (Fig. 5B).

**Figure 5.**
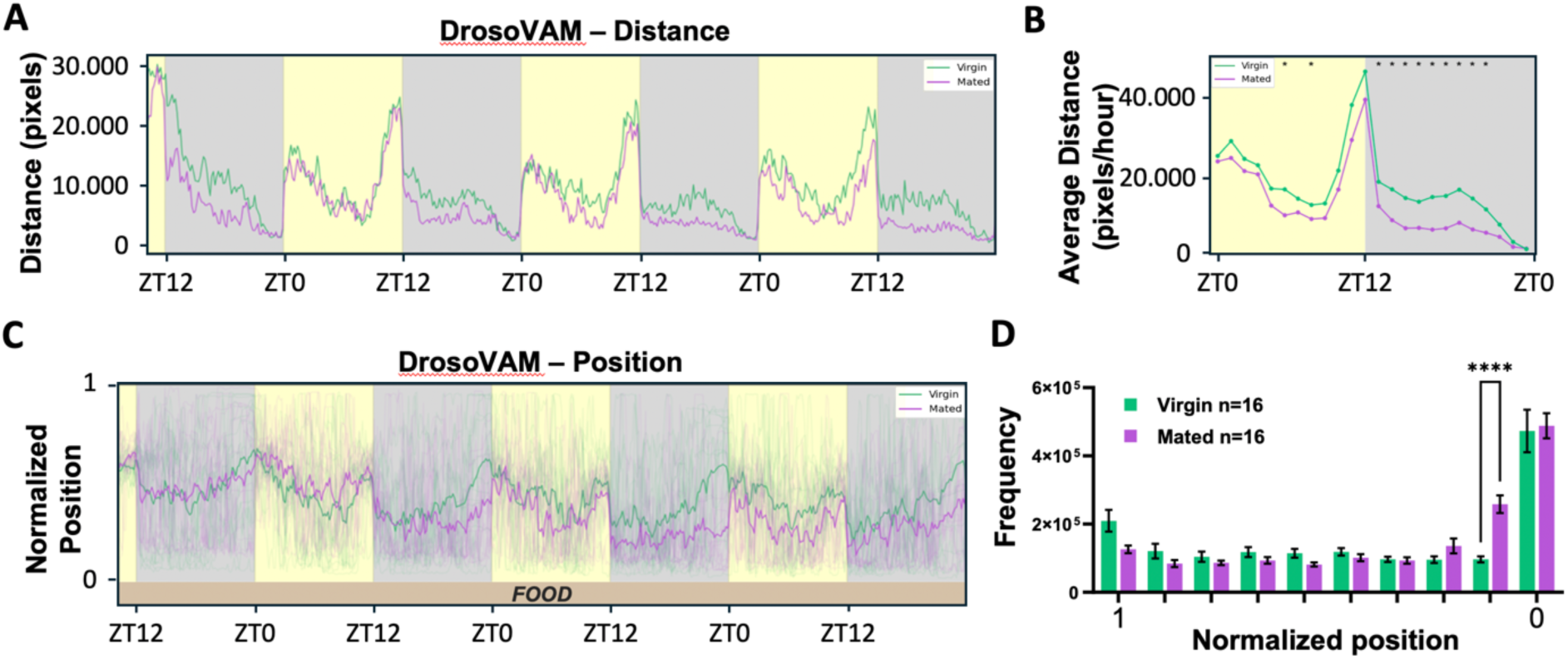
Analysis of the circadian activity of mated and virgin Drosophila hydei. Light/Dark periods are shown with yellow/grey background respectively. A. Distance in pixels moved by mated and virgin females for 86 hours post-male presentation, bins of 10 minutes, n=16/16. B. Average Day of the distance moved by mated and virgin female over 24 hours. Average day is composed of three days of monitoring. Statistical analysis was performed using tests of Mann-Whitney. C – Temporal representation of the normalized position of the flies in the locomotor activity chambers. The extremity of the chamber in which food is present is located at the bottom of the chart. Positions of all the flies are shown (transparent lines) and the average position is represented with the solid line, bins of 10 minutes. D – Histogram of the position of the flies in the DrosoVAM chamber over 86 hours. The side of the chamber where food is located is represented on the right of the histogram. Statistical analysis was performed using ANOVA2 (“Position X Mating Status” p-value < 0.0001), followed by Šídák’s multiple comparisons test (****: p-value < 0.0001).

Comparing virgin and mated locomotor activity over several days post-mating suggested an overall reduction of the activity in mated females (Fig. 5A-B). Interestingly, this reduction in movements was found to be more important at night, and to be maintained for more than 72 hours after mating. To understand why mated flies move less than virgins, we decided to look at the position of the flies. As DrosoVAM is based on video tracking, we could plot the position of flies over time relative to the food source. We identified that around 36 hours after mating, mated females start to stay closer to the food than their virgin counterparts (Fig. 5C). This effect could be confirmed by examining the overall position of the flies over the duration of the monitoring, showing that mated females are more commonly found on or near the food relative to virgin flies (Fig.5D). Interestingly, using the same methodology, we found that males also tend to stay further away from the food than females (Supp fig 5C).

### VI – Mating triggers a shift in light preference but not in food source

In *Drosophila melanogaster*, Sex Peptide has been found to trigger a shift in food source preference in mated females; mated females show a preference for yeast over sugar (32, 33). Such a food preference assay can be performed using DrosoVAM by examining fly positions in chambers with two food sources. For this assay, we tested mated female *D. hydei* in chambers with either an agar mix complemented with 2% yeast or 5% sugar, a condition that we previously showed to work for mated *D. melanogaster* females (Revel, submitted). However, when using this assay for *D. hydei*, we could not identify any food preference, in virgin or mated flies (Fig. 6A).

**Figure 6.**
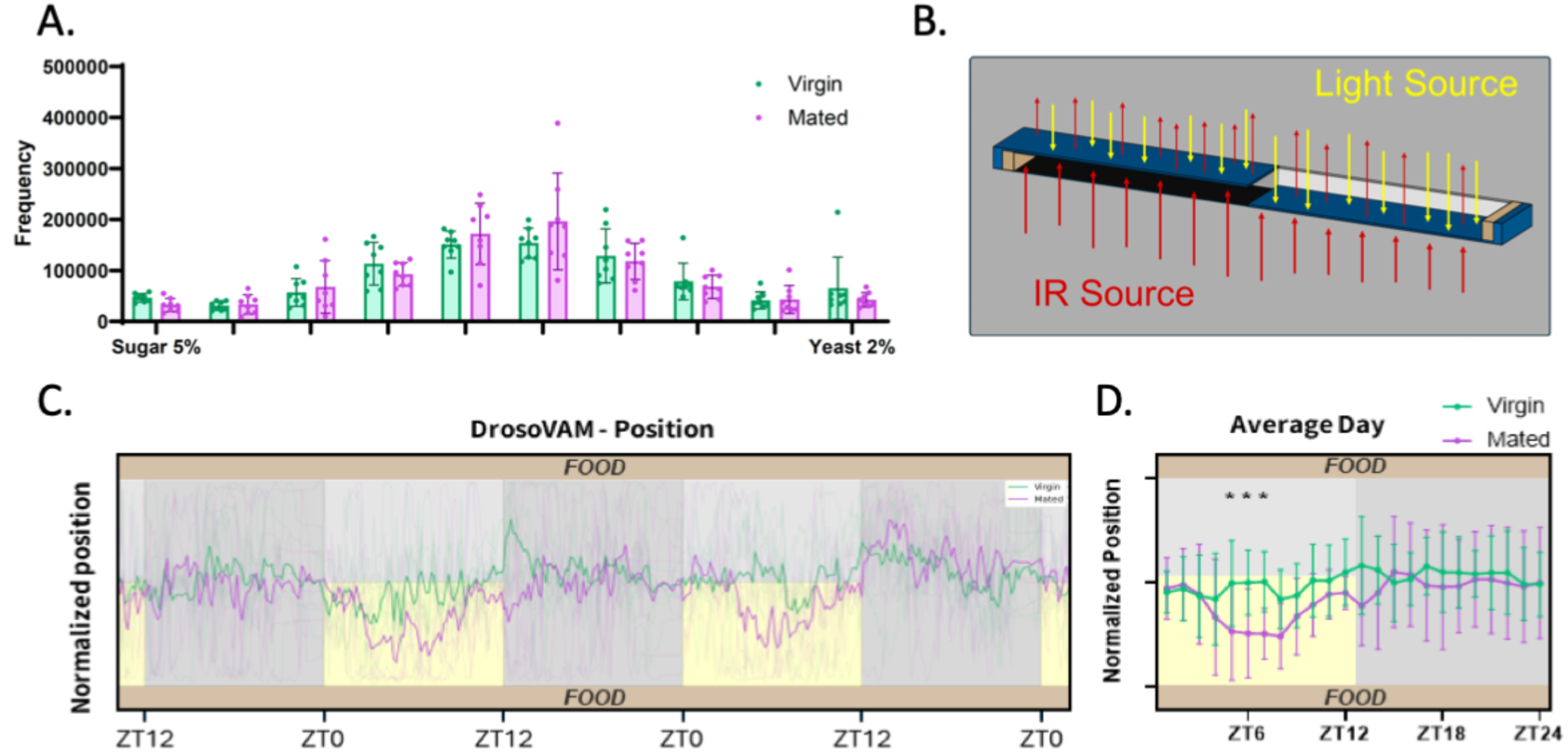
Position of the flies along the food/ light preference chambers. A. Histogram of the position of the flies along the food preference chambers. The chambers are divided into 10 bins (∼5mm per bin) and the amount of time each fly could be found in that position over 24 hours was quantified. The side of chamber with yeast-containing food is placed to the right while the side of the chamber with a simple sugar agar solution is placed to the left. T-tests performed between the positions closest to the food and between virgin and mated females (n=16/16) and found no significant difference. B. Design of a single Darkness Preference Chamber. The basic shape of the chamber is the same as for a food preference assay. There are visible light blocking/almost infrared transparent filters on both the right and left side of the chamber that are blocking visible light and are almost transparent to infrared, however the one pictured on the right is below the chamber relative to the visible light. This was necessary because of the slight reduction of the IR light transmitted to the camera (from below the chamber relative to the camera). Without the two filters, tracking quality decreased as contrast between the two sides of the chamber was vastly different. In these chambers, identical food sources are placed at both ends. C. Temporal representation of the position of the females in the darkness-preference assay chambers. Positions of all the flies are shown (transparent lines) and average position is represented with the solid line. D. Average position of flies at each hour of the day over the course of the experiment. Statistical analysis was performed using ANOVA2 ( “Time X Mating Status” p-value = 0.0211), followed by Šídák’s multiple comparisons test (p-values: ZT5 = 0.0477, ZT6 = 0.0294, ZT7 = 0.0216).

Because of the differences that we found between night and day localization of mated females, we wondered if the change in localization could be linked to light exposure. Taking advantage of the versatility of the DrosoVAM system, we decided to transform the food preference chambers into darkness preference chambers to evaluate if mated flies would change their light preference relative to virgins. For these assays, we shaded half of the chamber with a visible light opaque film-half of the chamber with identical food sources at both ends. Interestingly, in this assay, we observed a slight but significant preference of the mated females for staying in the non-shaded part of the chamber during lighted hours, whereas virgin flies displayed no preference (Fig.6 C-D). Meanwhile, it seems that males mostly prefer to avoid the light during the day (Supp fig 7). Although the purpose of this behavior which is still unclear, this phenotype is a novel PMR that may lead to a better understanding of the ecology of *D. hydei*.

## Discussion

### The egg-laying strategy of *D. hydei*

Although much is known about the reproductive strategies of the model species, *Drosophila melanogaster*, much less is known about other Drosophilids. This statement holds true for the characteristics of the female post-mating responses. In *D. melanogaster*, we know that the protein SP initiates a series of PMRs that cause the female to, among other things, produce and lay more eggs, change her diet and activity patterns, and be less receptive to remating. The gene for SP is relatively well conserved in the Drosophila genus but has been subject to gene duplication events in many species of the Sophophora subgenus (11), indicating that in these species, the SP response may have increased its relative importance.

It is often said that the PMRs were selected as strategies to maximize reproductive success. However, we have known for some time that many Drosophila species have lost the SP gene. As some of these species live in the same environments as *Drosophila melanogaster*, we wondered how the PMR responses have changed in these species and how these species initiate PMRs in the absence of SP. In this study, we have attempted to shed some light on the reproductive characteristics of the SP-less, cosmopolitan *Drosophila* species*, D. hydei*.

Unlike its more studied cousin, *Drosophila melanogaster*, we find that *D. hydei* does not seem to induce egg production after mating. Instead, we find that both virgin and mated females tend to lay a similar number of eggs over a three day period. What seems to change is that mated females will lay more eggs in the first 24 hours after mating, but then start laying less eggs than their virgin siblings, before reaching a virgin-like state again. This seems consistent with the idea that mating does not trigger an increase in oogenesis but instead triggers a short-term egg laying response to release stored eggs. Based on our daily counting of eggs from virgin females, we find that virgins do not lay the eggs at a constant daily rate, but instead seem to store them for a given period of time before “discarding” many eggs at one time. This strategy means that upon mating a female may occasionally have only a few eggs to lay but generally will have a stock of eggs to use instantly upon sperm transfer. One might conclude that such a strategy would be risky for the female, as they may be caught without a stock of eggs. However, this may be of minor importance in the wild if males are largely available and females are not at risk of struggling to mate. In this sense, females in the wild have been observed to remate up to four times with different males (34), suggesting that mates may be accessible enough so that females may benefit from a constant egg production, rather than waiting for egg production to be induced.

The difference in egg production strategy relative to *D. melanogaster* would be expected to have a number of consequences on the life of a *D. hydei* female. For example, *D. melanogaster* females change their dietary preference after mating, shifting to a diet higher in protein, thought to help sustain egg production. Indeed, preventing the dietary gut changes required to benefit from this change in diet has been shown to reduce egg-laying in mated *melanogaster* females (35). As *hydei* females do not experience an increase in egg production after mating, this type of dietary change would not be necessary. This hypothesis is consistent with our food preference assay results that show that mated and virgin *hydei* behave similarly when given a choice between two food options (sugar or yeast). Likewise, the increased metabolism required for egg production and/or the presence of SP has been linked to an overall decrease in female lifespan. As *hydei* females do not experience an increase in egg laying and do not have SP, one might expect females to live equally long in both virgin and mated conditions. Our survival assay shows that this is the case.

As mentioned above, we and others found that *hydei* females often mate more than once over a short window of time. These matings would generally last for around 2 minutes. We quantified the number of progeny resulting from both single and multiple matings and found that the immediate remating behavior that is observed in *D. hydei* has no impact on the number of eggs that are laid. This was interesting to us, as we initially thought that the short matings and the long sperm present in *hydei* would benefit from multiple matings to accumulate sperm in the female. As this does not seem to be the case and since it seems likely that female egg production limits the number of total progeny, we wondered if sperm competition might be a reason to have multiple matings. As we had limited tools in *hydei*, we attempted to test this by using male starvation as a proxy for male health status. We were able to show that male starvation seemed to affects male size, and the length and structure of their male accessory gland. However, the germline portion of the male reproductive tract was unchanged, at least with respect to testes length. As testes length correlates to sperm length, we believe that sperm length is probably unchanged by male starvation. This would be similar to egg length in the female, which we found does not change significantly between starved and unstarved females. We also checked if male starvation changed the number of sperm stored by the female by examining the number of progeny sired by starved or unstarved males. These assays also showed no significant difference. Thus, we conclude that starved males likely produce sperm of a similar length and deposit a similar number of sperm after a single mating. Interestingly, however, when we test the competitiveness of starved male sperm relative to unstarved male sperm, we find that sperm from starved males seems less competitive than sperm from unstarved males. Unlike melanogaster, we find that *hydei* often display a first male precedence, where after two sequential matings, more progeny are derived from the first male rather than the second. However, if the first male was starved its sperm seems less able to repel the sperm of the second male from being stored in the female.

At the present, we do not know why this is the case. One hypothesis is that the changes in the male accessory gland might allow females to judge the health of a male. As mentioned above, starvation seems to result in a reduction in overall MAG length and secondary cell number. This may lead to a quantitative change in the seminal fluid composition. In *D. melanogaster*, sperm competition has been linked to the protein SP that is attached to sperm via molecules largely produced by the secondary cells of the MAG. While *D. hydei* do not have SP, they actually have much larger number of secondary cells. Given the conservation of both secondary cells and the products associated with attaching SP to sperm in melanogaster, it is interesting to speculate that one of the secondary cell products might be linked to a mechanism through which females can use to judge male quality. If this is the case, then the selection for an increased number of secondary cells in many species might not be surprising, especially in the absence of SP.

### The circadian activity of *D. hydei*

Here, we also discovered a persistent behavioral switch in the circadian activity of mated female *D. hydei*. This change is characterized by reduced activity during night-cycles along with a preference for being in closer proximity to the food. The change in localization of the females upon mating is something that is also observed in *D. melanogaster*, though melanogaster does not share the drop in the night-time locomotor activity (31). Two reasons could possibly explain this preference to stay on/near the food: the need to lay eggs, and the need to eat. Egg laying has been found, in isolated *D. melanogaster* to be happening preferentially in the dark(36), which may explain the need to move closer to a food source at night. Interestingly, males seem to segregate away from the food source. Thus, another possibility for mated female relocation could be that this relocation is a way for the mated females to partition themselves away from males for one reason or another.

### The preference for light of mated females

In the wild, most *Drosophila* flies have been reported to be found in shaded areas during the daylight hours more than in full light (37). Thus, it is puzzling to find that mated females seem to preferentially move towards the light side of the chamber. We still do not understand why females display this behavior as one might guess that being in the lighted area would place them more at risk for predation. One hypothesis is that this could be a location-based mechanism to prevent the female from male interaction as males prefer to stay out of plain light. An alternative hypothesis is that our experimental setup might create a temperature difference on the two sides of the chamber; it is possible that the shaded side of the chamber is warmer due to the higher exposure to IR light due to slight IR blocking characteristics of the visible light filters (Fig. 6B). However, if temperature is the reason behind the behavior observed, it would mean that mated females prefer cooler areas during the day and not at night, as the IR light is on for the whole visible day/night cycle.

### How are the PMRs of *D. hydei* induced?

In *D. melanogaster*, many of the long-term PMRs are directly caused by the long-lived presence of SP in the reproductive tracts of the female. This is due to SP attaching to sperm and being stored in the sperm storage organs. Without being attached to sperm, most SP would likely be purged from the female along with the many sperm transferred by the male that fail to be stored. The absence of SP in *D. hydei* raises questions on the molecular mechanisms that may have evolved to cause the PMRs that we observe and how their effects might be maintained over time. The high number of Secondary Cells found in the MAG of *D. hydei* could suggest that their product plays a role in this process. Indeed, in *D. melanogaster*, Secondary Cells have been found to produce many of the proteins involved in the attachment of SP. Thus, if SC proteins of *D. hydei* are also capable of attaching to sperm, we might hypothesize that a mechanism could have arisen to use these proteins to bind other SFPs to sperm tails to induce the PMRs. Such a mechanism would create selective pressure to keep the SCs in the absence of SP, and maybe even provide enough pressure to increase the number of secondary cells. Interestingly, species in the *repleta/nannoptera* radiation also tend to have longer sperm tails than *Sophophora* species. *D. hydei*, for example, has sperms of 2.3cm (22), while *D. melanogaster* has sperms of less than 2mm (22). It is possible that, if SC proteins are capable of triggering PMRs in *D. hydei* and related species, and if these proteins still bind to sperm tails, it might actually favor males with more SCs and with longer sperm.

### The diversity in PMR strategies

Altogether, we believe that the absence of SP, a prominent inducer of PMRs, could open the door to the evolution of alternative ways for males to ensure their success over their rivals. Here, we have shown examples of different PMR in *D. hydei*, but there are many others. For example, *D. pegasea* and *D. mainlandi*, two other species without SP were found to have independently evolved a mate-guarding mechanism where the male is found riding the back of the female for hours after initial mating completion (38, 39). In a similar fashion, *D. acanthoptera,* which also lacks SP, was found to maintain mating for more than three hours in experiments conducted in the lab, which could be another form of a mate guarding strategy. These phenotypes, in essence, produce much the same effect as the SP-induced reduction in female receptivity.

The differences of the PMR observed in *D. hydei* compared to what is known in *D. melanogaster* suggests that many reproductive strategies may be equally valid in the wild. While both *melanogaster* and *hydei* are cosmopolitan species, differences in their ecological niche may have promoted the development of different mechanisms. Interestingly, this raises the question of the way reproductive biology is studied, as most experiments are carried out in the laboratory, where conditions are simplified to the extreme. Fundamentally, observations that we made regarding the post-mating behavior of *D. hydei* cannot be explained at this time and require deeper investigation of both the molecular mechanisms and ecological forces that are at play.

## Material and Methods

### Drosophila strains

*Drosophila hydei* strains were kindly provided by Benjamin Loppin (LBMC, ENS, Lyon, France), all other non-*melanogaster* species were kindly provided by Markus Knaden (nGICE, Max Planck Institute, Jena, Germany). All species were dissected at reproductive maturity upon arrival and/or housed in standard conditions at 25°C on standard *Drosophila melanogaster* cornmeal food.

### Dissection and imaging

#### Accessory Glands

Males of each species are isolated from same-age stocks a few days after reaching sexual maturity and dissected in 0.3% PBS-Triton. Glands are isolated and transferred into 1 ml of 0.3% Triton-PBS. Glands are fixed by adding 110 µl of 36% Paraformaldehyde and incubating at room temperature for 10 minutes on rotating wheel. Without removing the PFA, glands are then stained by adding 120 µl of Ethidium Bromide at 1% and incubating for another 10 minutes at room temperature on a rotating wheel. All liquid is removed and the glands are washed twice for 10 minutes in 1 ml 0.3% PBS-Triton. Glands are then transferred onto a slide with Vectashield, covered with a cover slip and sealed with nail polish. Glands were imaged using the Ni-E confocal microscope from Nikon with the NIS-Element C imaging software. Images were then analyzed using Fiji (ImageJ2) to measure the glands, and Secondary cells are counted using the Cell Counter Fiji plugin (https://imagej.net/ij/plugins/cell-counter.html).

#### Eggs

The eggs of the flies that were used in the malnourished vs nourished egg-laying assay were collected from the tubes and separated from the food in 1X PBS. The clean eggs were transferred onto a slide and fixed with 3.6% Paraformaldehyde. The slides were covered with a cover slip and sealed with nail polish. Imaging was done using Axioplan2 2 and Fiji (ImageJ2) was used to measure the length

#### Thorax and testes size

*D. hydei* males that were used in the sperm competition assays were dissected in 1X PBS. Their thorax and testes were isolated and transferred onto a slide and then fixed with 3.6% Paraformaldehyde. Thoraces were placed into a channel made by gluing stacks of coverslips to a slide. Space within the channel was filled with 10%glycerol and the whole area was covered with an additional coverslip and sealed. Testes were dissected and extended on slides in PBS, in a way that made measuring possible while trying to minimize tissue stretching. We measure testes in a way that included the seminal vesicle due to the difficulty in judging exactly where each part began/ended. The slides were covered with a cover slip and sealed with nail polish. Imaging was done using Leica DM5500 B. Fiji (ImageJ2) was used to measure the size.

### Mating and egg laying assays

Each female was individualized a few days prior to mating and placed on a 12:12-hour light:dark cycle. About 1 hour before expected lights-on, ∼10-day old male *Drosophila hydei* of the appropriate genotype were added to each tube (diameter 13 mm) and visually monitored until mating occurred. Upon mating completion, the male was either allowed to remate with the female, in the case of multi-mating assays, or removed. For egg laying assays, virgin or mated females were then transferred to new tubes and left at 25°C. Tubes were then changed every 24 hours and the number of eggs were counted. Tubes were then kept to verify the fertilization of the eggs. Egg numbers were then entered into GraphPad PRISM (Version 10) for subsequent analysis and plotting.

### Sperm competitions

For sperm competition assays, a similar mating protocol was used as above. For these assays, w- female flies were used and mated to either w- or w+ flies depending on the experiment. We performed the experiment in both directions and found that there was no change linked to the first male being w- or white+. Also, due to the overall lower mating rate in hydei, we performed these experiments over multiple days and combined all of the results.

In order to check for the influence of nutritional state on sperm competitiveness, we placed ∼70-100 embryos from overnight egg layings of w+ or w- stocks onto tubes containing either 8 ml normal food (per 10L standard food: 0.65 % (w/v) agar, 6% (w/v) powdered yeast, , 6% (w/v) corn meal, 7% (w/v) sugar, 0.3% (w/v) Nipagine, 1.7 % (v/v) propionic acid) or an equal volume of normal food diluted 1:3 with 1% agar. Males were collected and aged in groups of 20-25 males for ∼10 days in normal food.

Mating was performed as above. Random males that mated were measured for their thorax size and their testes length (see above).

Female progeny were counted and scored for eye color. Due to occasional false mating, only tubes with both w- and w+ progeny were counted as having mated multiple matings.

### Survival assay

For the intermittent mating survival assay a total of 80 virgin females placed in tubes in groups of eight or nine flies per tube. The 89 males were also divided into tubes in groups of eight or nine. At Day 6, an equal number of 10 day old males were added to each 5 tubes, with the remaining tubes were kept as virgins. Overall 44 females were mated. And 36 females (less due to some flies escaping) were kept as virgins. Both a week and two weeks later, the matings were repeated. Flies were transferred every 2-3 days into fresh tubes, each time counting dead flies and removing them.

For a constant mating survival assay, 100 males and 96 females were kept as virgins, while 49 females were housed with 50 males. The flies were divided into tubes in groups of 20 of the same sex, and or approximately 10 females and 10 males. Tubes were changed every two to three days as above, and dead flies were counted.

### Locomotor activity, position analysis and preference assays

Males, virgin females, or mated females were loaded into 3D-printed chambers, recorded, tracked and analyzed as described in (31). To set up the darkness-preference assay, the food preference arena has been modified using an IR-pass plastic. Such plastic was obtained using the magnetic band of old floppy disks (diskette). IR-pass plastic was applied differentially on the upper- and lower-halves of the chambers. On the upper part, plastic was applied on the top of the chamber, blocking most of the visible light, and dimming only slightly IR light. The plastic was also applied on the bottom of the lower half of the chamber to not block any visible light, while still equally dimming IR light. The interface between the two halves of the chamber caused a slight alteration in the ability of DeepLabCut (40) to detect flies crossing the interface but nevertheless allowed us to track the flies in the rest of the chambers.

---

**Supplementary Figure 1:**
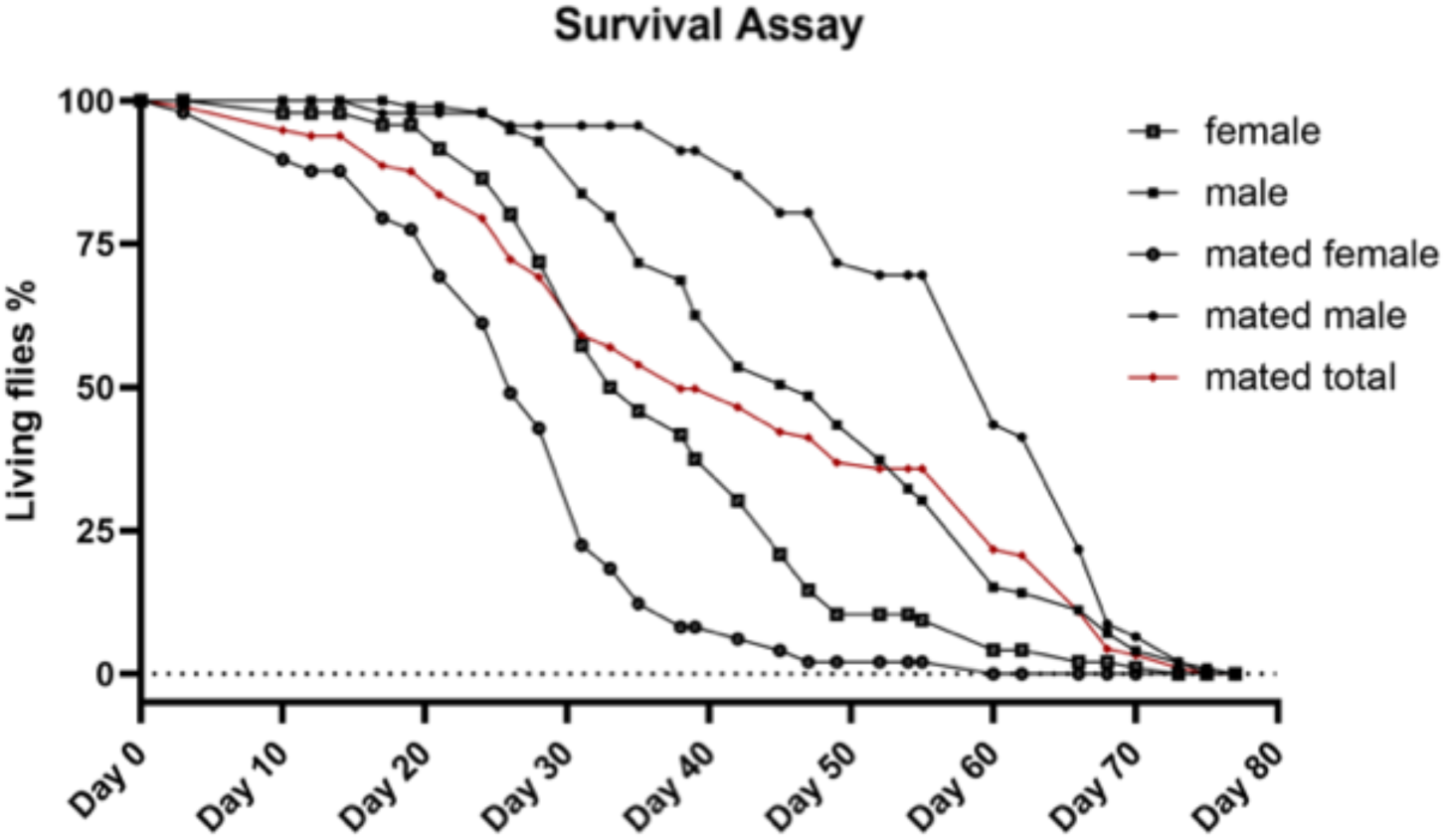
Survival assay where mated females and mated males were housed together.

**Supplementary Figure 2:**
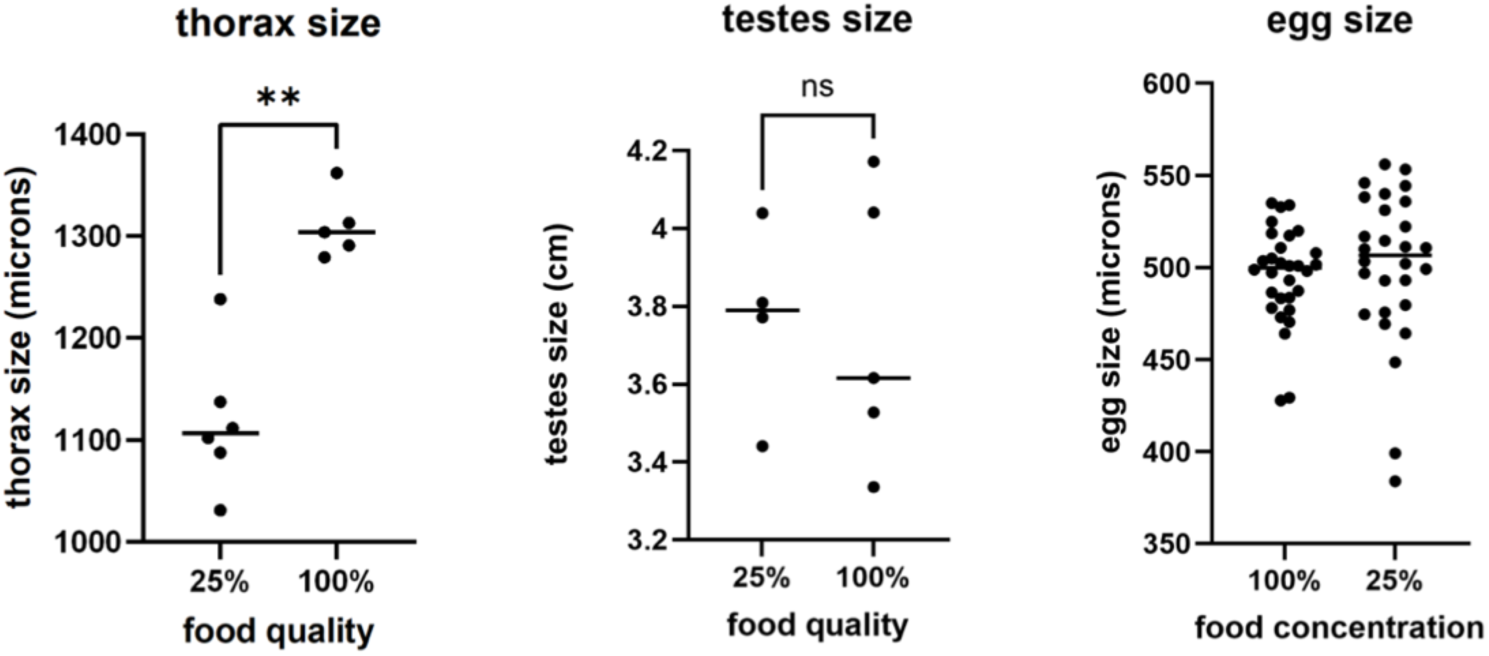
Characteristics of flies after growth on lower quality food. Non-parametric t-test were performed on each characteristic measured. Only thorax size was significantly affected by these manipulations.

**Supp Fig 3.**
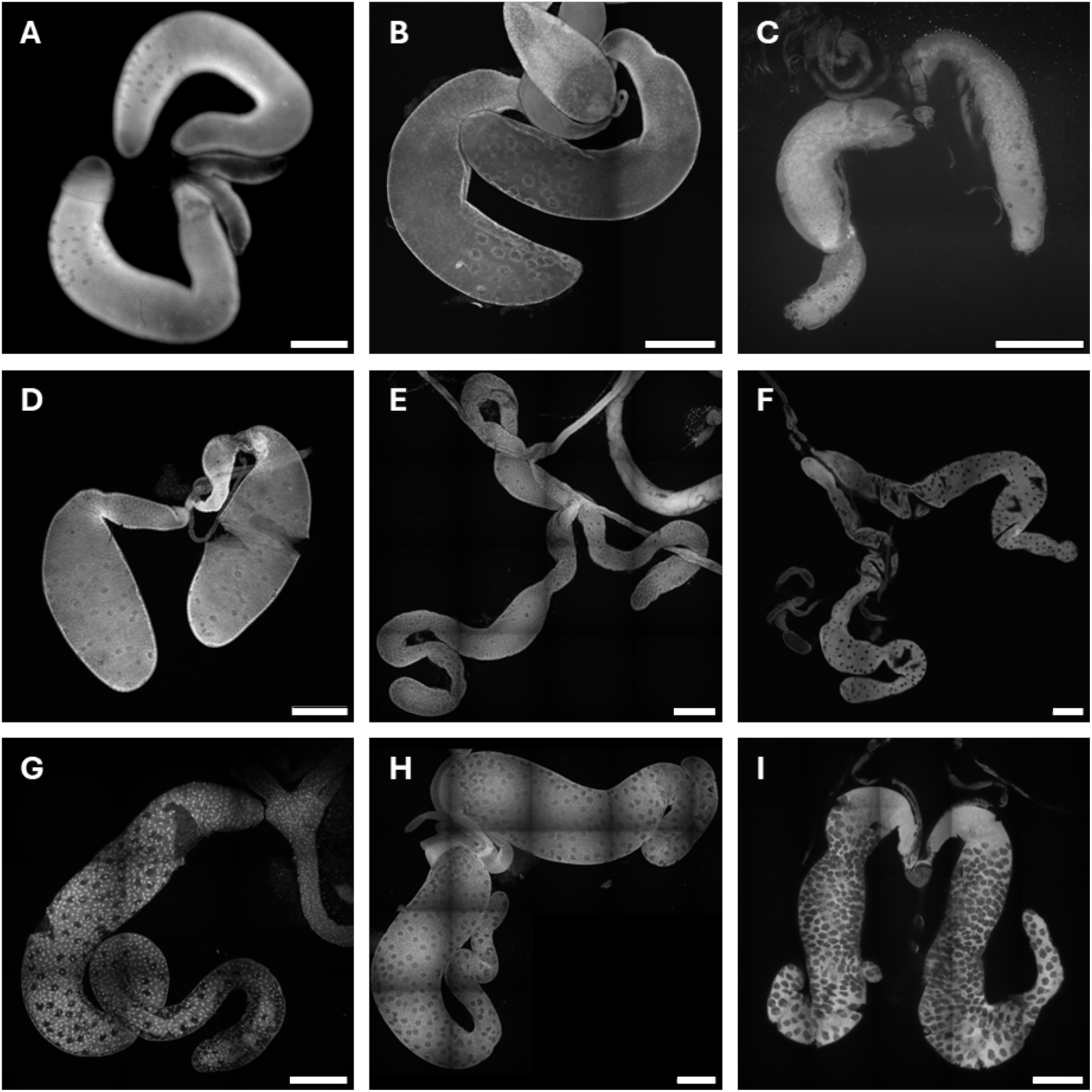
Accessory Glands of several Drosophila species. A – *Scaptodrosphila lebanonensis*; B – *Drosophila melanogaster*; C – *Drosophila greeni*; D – *Drosophila ananassae*; E – *Drosophila virilis*; F – *Zaprionus indianus*; G – *Drosophila bifurca*; H – *Drosophila hydei*; I – *Drosophila nannoptera*. Ethidium Bromide staining, scale bars represent 200µm.

**Supplementary Figure 4.**
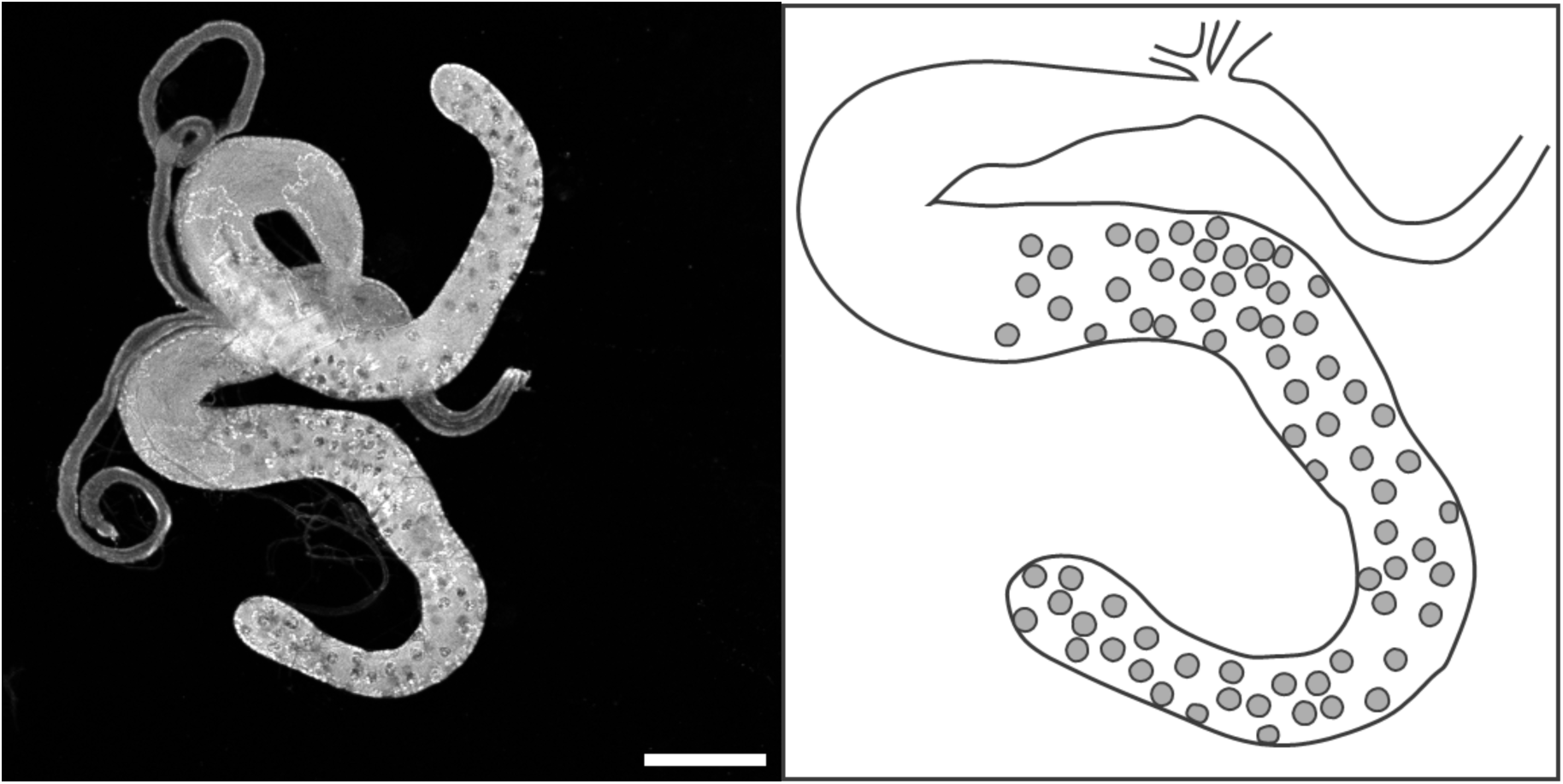
Left. Accessory glands of Chymomyza pararufithorax. Ethidium Bromide staining, scale bars represent 200µm. Right. Representation of a gland lobe, with the secondary cells spread along most of the lobe length.

**Supplementary Figure 5.**
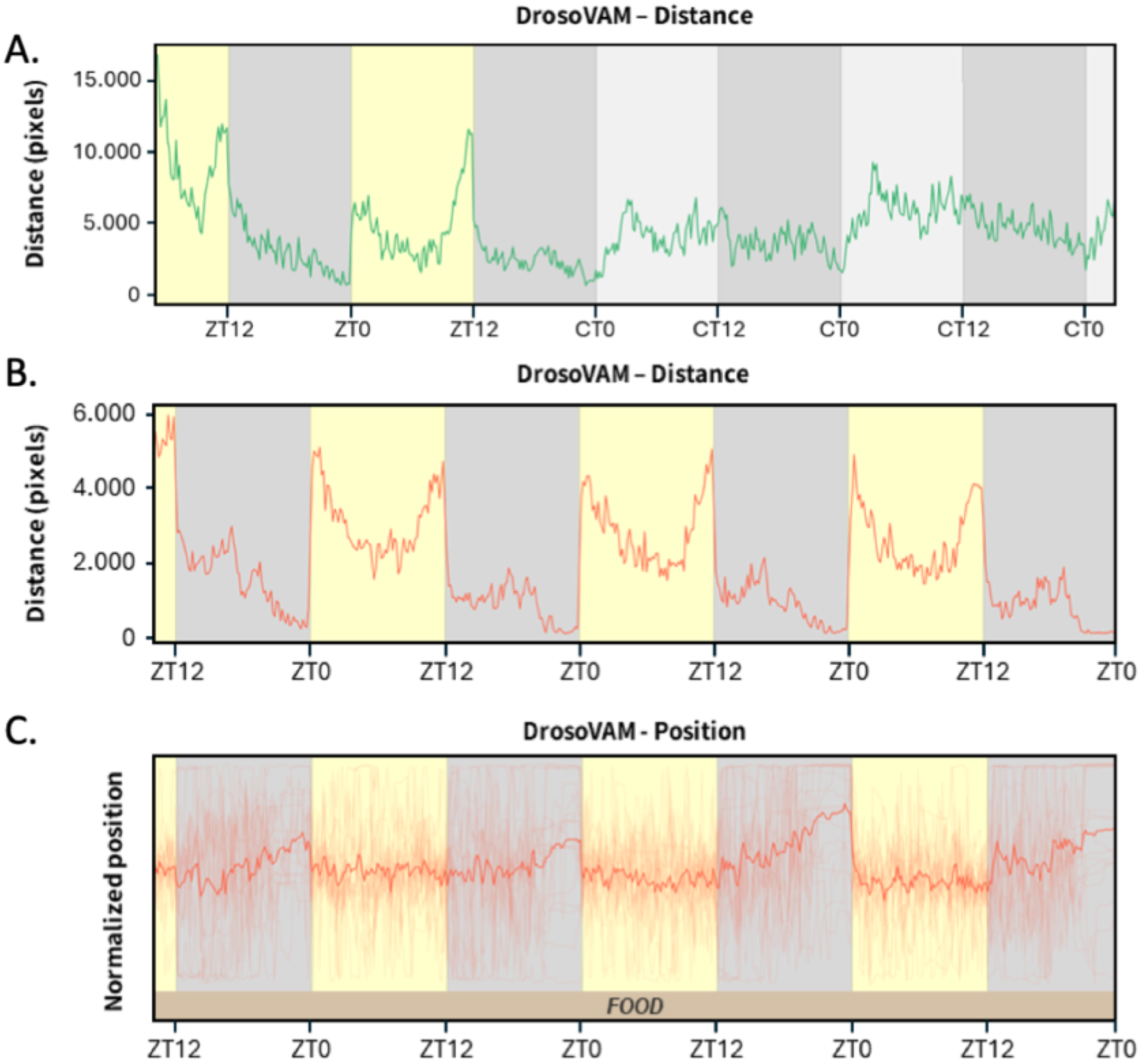
Further analysis of the circadian activity of Drosophila hydei. A. Flies were trained in 12h:12h Light/Dark cycles and then switched to constant darkness. Light/Dark cycles are shown with yellow/grey background respectively. In constant darkness, subjective day/night periods are shown with light grey/ grey background. Distance in pixels moved by virgin females, bins of 10 minutes. Circadian variations in the position relative to food of males *Drosophila hydei.* B. Distance in pixels moved by males. C. Temporal representation of the position of the flies in the locomotor activity chambers. The extremity of the chamber in which food is present is located at the bottom of the chart. Positions of all the flies are shown (transparent lines) and the average position is represented with the solid line.

Supplementary Figure 6

**Supplementary Figure 7:**
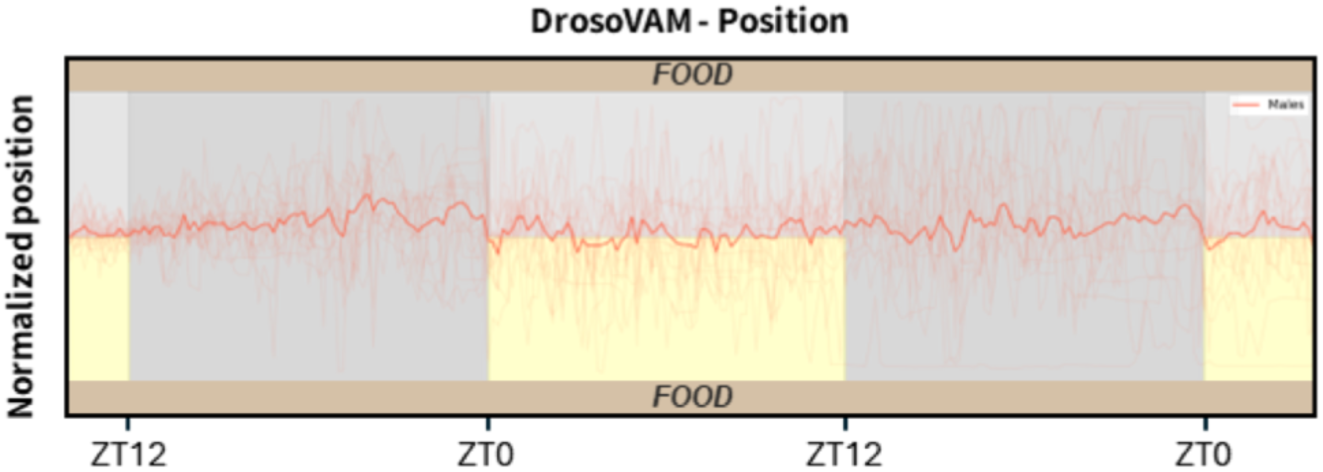
Temporal representation of the position of the males in the darkness-preference assay chambers. Positions of all the flies are shown (transparent lines) and average position is represented with the solid line.

## Notes

### Competing Interest Statement

The authors have declared no competing interest.

### Funding

- Swiss National Science Foundation: 310030_192621, 320030-227485
- Canton of Geneva
- University of Geneva
- George and Antoine Claraz Foundation

